# Lorcaserin maintenance fails to attenuate heroin vs. food choice in rhesus monkeys

**DOI:** 10.1101/705020

**Authors:** E. Andrew Townsend, S. Stevens Negus, Justin L. Poklis, Matthew L. Banks

## Abstract

**Background:** The current opioid crisis has reinvigorated preclinical research in the evaluation of non-opioid candidate treatments for opioid use disorder (OUD). Emerging evidence suggests 5-HT_2C_ receptor agonists may attenuate the abuse-related effects of opioids. This study evaluated effectiveness of 7-day treatment with the clinically available 5-HT_2C_ agonist lorcaserin on heroin-vs.-food choice in rhesus monkeys. Lorcaserin effects were compared to effects produced by saline substitution and by 7-day treatment with the opioid antagonist naltrexone.

**Methods:** Adult male (1) and female (6) rhesus monkeys were trained to respond under a concurrent schedule of food delivery (1g pellets, fixed-ratio 100 schedule) and intravenous heroin injections (0-0.032 mg/kg/injection, fixed-ratio 10 schedule) during daily 2h sessions. Heroin choice dose-effect functions were determined daily before and following 7-day saline substitution or 7-day continuous treatment with naltrexone (0.0032-0.032 mg/kg/h, IV) or lorcaserin (0.032-0.32 mg/kg/h, IV).

**Results:** Under baseline conditions, increasing heroin doses maintained a dose-dependent increase in heroin choice. Both saline substitution and 7-day naltrexone treatment significantly attenuated heroin choice and produced a reciprocal increase in food choice. Continuous lorcaserin treatment significantly increased heroin choice.

**Conclusions:** In contrast to saline substitution and naltrexone, lorcaserin treatment was ineffective to reduce heroin-vs.-food choice. These preclinical results do not support the therapeutic potential of lorcaserin as a candidate OUD treatment.

## 1.0 Introduction

Despite the availability of methadone, buprenorphine, and naltrexone as Food and Drug Administration (FDA)-approved pharmacotherapies for opioid use disorder (OUD) in the United States, the opioid crisis remains a significant public health issue. One reason for the ongoing opioid crisis is that the majority of patients who meet diagnostic criteria for OUD do not seek medical help (Blanco et al., 2013; Han et al., 2018). Other reasons include poor treatment adherence (especially for naltrexone), barriers to treatment access (especially for methadone), and abuse-liability concerns (for methadone and buprenorphine) (Blanco and Volkow, 2019; Volkow et al., 2014). Furthermore, a proportion of patients who do receive treatment will still relapse, highlighting the clinical need for other OUD monotherapies or adjuncts to existing FDA-approved treatments (Lee et al., 2018). In response to the opioid crisis, the National Institutes of Health have outlined several strategies to focus scientific efforts culminating in the “Helping to End Addiction Long-Term” (HEAL) initiative (Collins et al., 2018; Volkow and Collins, 2017). One prioritized area for preclinical research is the development and evaluation of novel OUD medications.

The abuse-related effects of mu opioid receptor (MOR) agonists are thought to be mediated at least in part by activation of mesolimbic dopamine (DA) neurons that originate in the ventral tegmental area and project to forebrain regions such as the nucleus accumbens (Koob and Volkow, 2016). More specifically, mesolimbic DA neurons receive inhibitory inputs from GABA-ergic interneurons that express G_i/o_-protein-coupled MORs, and MOR agonist binding to these receptors inhibits GABA neuron activity to disinhibit and activate downstream mesolimbic DA neurons. These GABA-ergic neurons also express excitatory G_q_/G_11_-protein-coupled 5-HT_2C_ receptors (Hannon and Hoyer, 2008), and 5-HT_2C_ agonists have the potential to oppose and limit MOR agonist effects on activity of both GABA-ergic neurons and downstream mesolimbic DA neurons. For example, acute pretreatment with the 5-HT_2C_ receptor agonist MK-212 blunted morphine-induced increases in nucleus accumbens extracellular DA levels (Willins and Meltzer, 1998). Consistent with this neurochemical evidence, acute pretreatment with the 5-HT_2C_ receptor agonist lorcaserin decreased reinstatement of operant-responding previously maintained by remifentanil in rhesus monkeys (Gerak et al., 2019) or by oxycodone in rats (Neelakantan et al., 2017). Acute lorcaserin pretreatment also decreased oxycodone self-administration in rats (Neelakantan et al., 2017) and heroin self-administration in rhesus monkeys (Kohut and Bergman, 2018). Furthermore, lorcaserin treatment effects were both sustained during repeated lorcaserin administration and selective for heroin vs. food-maintained responding in rhesus monkeys (Kohut and Bergman, 2018). On the basis of these and related findings, the 5-HT_2C_ receptor has emerged as a pharmacological target of interest by the National Institute on Drug Abuse to address the opioid crisis (Rasmussen et al., 2019; Volkow and Collins, 2017).

To further evaluate 5-HT_2C_ agonists as candidate non-opioid treatments for OUD, the present study evaluated lorcaserin effectiveness to reduce intravenous (IV) heroin self-administration under a heroin-vs.-food choice procedure in rhesus monkeys. A preclinical heroin-vs.-food choice procedure was utilized for two reasons. First, although 5-HT_2C_ agonists can selectively reduce opioid self-administration more than food-maintained responding under some conditions, the magnitude of this selectivity appears to be modest (≤3-fold), and lorcaserin is clearly effective to decrease rates of food-maintained responding at doses similar to or only slightly higher than those that decrease rates of opioid self-administration (e.g. (Banks and Negus, 2017a)). Drug-vs.-food choice procedures provide an alternative strategy to dissociate selective treatment effects on opioid reinforcement (evaluated as changes in behavioral allocation between drug and food choice) from non-selective treatment effects on operant responding (evaluated as changes in overall behavioral rates maintained by either drug or food choice) (Griffiths et al., 1976; Hart et al., 2000; Johanson, 1975; Negus, 2006; Woolverton and Balster, 1981). Second, these procedures have increasingly demonstrated translational concordance with both human laboratory studies and clinical trials across a broad range of abused drug classes, including opioids (for review, see (Banks, 2017; Banks et al., 2015; Banks and Negus, 2017b; Negus and Banks, 2013)). To provide a framework for interpreting lorcaserin effects, the effects of two other manipulations were examined. First, saline was substituted for heroin for seven consecutive days to examine extinction of heroin choice when heroin was absent, but the discriminative stimuli were still present. Second, monkeys were treated continuously for seven consecutive days with each of three different doses of naltrexone (0.0032-0.032 mg/kg/h). Naltrexone served as a positive control because depot naltrexone formulations are FDA-approved for OUD treatment in non-dependent patients (Comer et al., 2006; Krupitsky et al., 2011; Tanum et al., 2017). Human laboratory studies suggest that clinically effective depot naltrexone reduces opioid potency at least 8-fold (Sullivan et al., 2006), and any candidate OUD treatment should be at least as effective as these FDA-approved naltrexone formulations to warrant further consideration.

## 2.0 Methods

### 2.1 Subjects

Studies were conducted in a total of seven adult rhesus monkeys (*Macaca mulatta*) of Indian origin (one male, six females). Monkeys were surgically implanted with a double-lumen venous catheter (extruded silicone i.d. = 0.76 mm, o.d. = 2.36 mm, durometer = 60; Reiss Manufacturing, Blackstone, VA) for studies of heroin-vs.-food choice. Three female monkeys had prior opioid drug discrimination histories, the male monkey had prior cocaine self-administration history, and the other three female monkeys were experimentally naïve. Monkeys could earn 1-g banana-flavored pellets (5TUR Grain-based Precision Primate Tablets; Test Diets, Richmond, IL) during daily experimental sessions (see below). The monkeys’ diet consisted of food biscuits (Teklad Global Diet, 2050, 20% Protein Primate Diet) and fresh fruit or vegetables delivered in the afternoons after behavioral sessions to minimize the effects of biscuit availability and consumption on food-maintained operant responding. Water was continuously available in each monkey’s home chamber, which also served as the experimental chamber. A 12-h light/dark cycle was in effect (lights on from 0600 to 1800 h). Environmental enrichment consisting of movies displayed on a monitor in the housing room and foraging devices loaded with nuts, seeds, or diced vegetables, was also provided after behavioral sessions. Facilities were licensed by the United States Department of Agriculture and accredited by the AAALAC International. The Institutional Animal Care and Use Committee approved all experimental and enrichment protocols. Animal research and husbandry were conducted according to the 8^th^ edition of the Guide for the Care and Use of Laboratory Animals.

### 2.2 Heroin vs. food choice procedure

Each housing chamber was equipped with a customized operant response panel, which had two response keys that could be transilluminated red or yellow, and a pellet dispenser (Med Associates, ENV-203–1000, St. Albans, VT) that delivered food pellets to a receptacle below the operant panel. The externalized portion of the intravenous (IV) catheter was routed through a custom jacket and tether system connected to a dual-channel fluid swivel (Lomir Biomedical, Malone, NY) on the chamber top and connected to two safety syringe pumps (Med Associates, PHM-108), one for each lumen of the double-lumen catheter. One pump (the “self-administration” pump) was used to deliver contingent IV heroin or saline injections through one lumen of the double-lumen catheter. The second pump (the “treatment” pump) was used to deliver non-contingent IV saline, naltrexone (0.0032-0.032 mg/kg/h), or lorcaserin (0.032-0.32 mg/kg/h) injections through the second lumen at a programmed rate of 0.1 mL injections every 20 min from 1200 h each day until 1100 h the next morning. Catheter patency was periodically evaluated with IV ketamine (Vedco, St. Joseph, MO) administration and after any treatment that produced a rightward shift in the heroin choice dose-effect function. The catheter was considered patent if IV ketamine administration produced muscle tone loss within 10 s.

Daily experimental sessions were conducted from 0900 to 1100 h in each monkey’s home chamber as described previously for heroin-, cocaine-, and methamphetamine-vs.-food choice studies (Banks and Blough, 2015; Banks et al., 2011; Negus, 2006). The terminal choice schedule consisted of five 20-min components separated by 5-min inter-component intervals during which responding had no scheduled consequences. During each component, the left, food-associated key was transilluminated red, and completion of the FR requirement (FR100) resulted in food pellet delivery. In addition, the right, heroin-associated key was transilluminated yellow, and completion of the FR requirement (FR10) resulted in delivery of the unit heroin dose available during that component. These response requirements were based on previously published studies wherein monkeys readily switched from the food-associated key to the heroin-associated key when an intermediate unit heroin dose was available (Negus, 2006; Negus and Rice, 2008). These response requirements were intended to position the heroin choice dose-effect function such that the detection of leftward and rightward shifts were possible following experimental manipulations (see (Banks and Blough, 2015; Negus, 2003)). The unit heroin doses available during each of the five successive components were 0 (i.e., no heroin), 0.001, 0.0032, 0.01, and 0.032mg/kg/injection, respectively, and dose was varied by changing the resulting volume (0, 0.03, 0.1, 0.3, and 1.0 ml/injection, respectively) of each injection. Stimulus lights on the heroin-associated key were flashed on and off in 3 s cycles, and longer flashes were associated with larger heroin doses. Ratio requirement completion initiated a 3 s timeout, during which all stimulus lights were turned off, and responding had no scheduled consequences. Choice behavior was considered to be stable when the lowest unit heroin dose maintaining at least 80% heroin choice varied by ≤0.5 log units for three consecutive days. Data from these three days were subsequently used as the “baseline” for statistical and graphical comparisons to each treatment.

Once heroin-vs.-food choice was stable, experimental test periods were conducted to determine effects of saline substitution and of treatment with naltrexone or lorcaserin on heroin-vs.-food choice. For saline substitution, saline replaced heroin in the self-administration pump for seven consecutive days and saline continued to be infused via the treatment pump. For naltrexone and lorcaserin treatments, heroin was retained in the self-administration pump, naltrexone or lorcaserin solutions replaced saline in the treatment pump, and each naltrexone (0.0032-0.032 mg/kg/h, IV) and lorcaserin (0.032-0.32 mg/kg/h, IV) dose was tested for seven consecutive days. Continuous infusion was selected as the route of administration for naltrexone and lorcaserin to allow for a greater opportunity for any tolerance to treatment effects to emerge and lessen concerns regarding whether the behavioral session coincided with steady plasma drug concentrations. For example, we have previously reported acute intramuscular lorcaserin effects on food-maintained responding were dissipating by the 135 min time point (Banks and Negus, 2017a), a time point that would correspond to the later components of the heroin-vs.-food choice procedure. The intravenous route was chosen to deliver the chronic infusion because venous access was already available through the implantation of the double-lumen catheter. At the conclusion of each 7-day saline-substitution or drug-treatment test period, heroin and saline solutions were reinstated as necessary in the self-administration and treatment pumps, respectively, and responding was monitored for at least 4 days and until the lowest unit heroin dose maintaining at least 80% heroin choice varied by ≤0.5 log units for three consecutive days. Following saline substitution or a drug treatment wherein a rightward or downward shift of the heroin-choice function was observed, the food-associated light was terminated in the last two response components for a single session, promoting behavioral allocation to the heroin-associated key. Subsequently, heroin-vs.-food choice sessions during saline maintenance were conducted until the aforementioned stability criteria were met. Each treatment was evaluated in at least four monkeys. Saline substitution was evaluated in all monkeys before drug treatments to ensure that seven days were sufficient to observe a decrease in choice of the heroin-associated key in the absence of heroin reinforcement. Five monkeys received both naltrexone and lorcaserin. The dose order within each drug was counterbalanced across subjects, and in those monkeys that received both naltrexone and lorcaserin, naltrexone was tested first because naltrexone was the positive control compound.

### 2.3 Naltrexone and lorcaserin plasma sample analysis

Blood was collected from monkeys under ketamine anesthesia at day seven during continuous naltrexone or lorcaserin treatment into vacutainer tubes. Tubes were immediately centrifuged at 1000g for 10 min, and the plasma supernatant was transferred into a storage tube and frozen at –80°C until analyzed. A seven-point calibration curve (naltrexone: 1-500 ng/mL, lorcaserin 10-1000 ng/ml) was determined, which included a drug-free control and a negative control without internal standard (ISTD) in drug-free plasma. Naltrexone was extracted from the plasma using a liquid/liquid extraction. In brief, 100 ng/mL naltrexone-d_3_ and the ISTDs were added to 100 µL aliquots of plasma of each calibrator, control, or specimen except the negative control. Next, 0.5 mL of saturated carbonate/ bicarbonate buffer (1:1, pH 9.5) and 1.0 mL of chloroform:2-propanol (9:1) were added. Naltrexone samples were mixed for 5 min and centrifuged at 2500 rpm for 5 min. The top aqueous layer was aspirated, and the organic layer was transferred to a clean test tube and evaporated to dryness at 40°C under a constant stream nitrogen. The samples were reconstituted with methanol and placed in auto-sampler vials for analysis. Lorcaserin was extracted from the plasma using the addition of acetonitrile. In brief, the ISTD, 10 ng of cocaine-d3, was added to aliquots of 100 µL of the calibrators, controls and samples. These samples were mixed and allowed to equilibrate. 0.2 mL acetonitrile was added to each sample and vortex mixed. The samples were then centrifuged at 2054 g for 10 min. After centrifuging the top layer containing the acetonitrile was removed and placed in auto-sampler vials for analysis.

The ultra-performance liquid chromatography tandem mass spectrometer (UPLC-MS/MS) analysis was performed for naltrexone and lorcaserin on a Sciex 6500 QTRAP system with an IonDrive Turbo V source for TurbolonSpray® (Sciex, Ontario, Canada) attached to a Shimadzu UPLC system (Kyoto, Japan) controlled by Analyst software (Sciex, Ontario, Canada). Chromatographic separation of naltrexone was performed using a Zorbax XDB-C18 4.6 x 75 mm, 3.5-micron column (Agilent Technologies, Santa Clara, CA). The mobile phase contained water:methanol (80:20, v/v) with 1 g/L ammonium formate and 0.1 % formic acid and was delivered at a flow rate of 1 mL/min. The source temperature was set at 650°C. Curtain gas flow rate was set to 30 mL/min. The ionspray voltage was 5500 V, with the ion source gases 1 and 2 having flow rates of 60 mL/min. The declustering potential and collection energy were 86 eV and 39 eV, respectively. The quantification and qualifying transition ions were monitored in positive multiple reaction monitoring mode: Naltrexone 342> 267 and 342 > 282 and Naltrexone-d_3_ 345> 270 and 345 > 285. For lorcaserin, chromatographic separation of lorcaserin and the ISTD, cocaine-d3, was performed using a Thermo Hypersil Gold column, 50 x 2.1 mm, 3 micron (Thermofisher Scientific, USA). The mobile phase contained water/methanol (40:60, v/v) with 0.1 mM ammonium formate and was delivered at a flow rate of 1 mL/min. The source temperature was set at 600°C, and curtain gas had a flow rate of 30 mL/min. The ionspray voltage was 5500 V, with the ion source gases 1 and 2 having flow rates of 60 and 45 mL/min, respectively. The declustering potential and collection energy were for lorcaserin and the ISTD were 100 and 38 V, respectively. The acquisition mode used was multiple reaction monitoring. The following transition ions (m/z) with their corresponding collision energy in parentheses were monitored in negative mode for lorcaserin: 196>144 (34) and 196>129 (40) and in positive mode for cocaine-d3: 307>185 (28) and 307>105 (50). The total run time for the analytical method was 4 min. A linear regression of the peak area of ratios of the quantification transition ions and the ISTDs transition ions were used to construct the calibration curves.

### 2.4 Data analysis

Two primary dependent measures were determined for each component: (1) percent heroin choice, defined as (number of ratio requirements, or ‘choices,’ completed on the heroin-associated key/total number of ratio requirements completed on both the heroin- and food-associated keys)*100, and (2) the number of choices completed per component. Mean data from the three days preceding each pharmacological treatment were averaged for each individual monkey and then averaged across monkeys to yield group mean “baseline” data. Mean data from the last 3 days of each 7-day naltrexone or lorcaserin were averaged for each individual monkey and then averaged across monkeys to yield group mean data. Percent heroin choice and number of choices completed per component were then plotted as a function of the unit heroin dose and analyzed using a repeated-measures two-way ANOVA (saline substitution, naltrexone) or mixed-effects analysis (lorcaserin) with treatment drug dose and unit heroin dose as the fixed main effects (Prism 8.0.1 for macOS, GraphPad Software, San Diego, CA). Additional dependent measures collected during each session were total choices, total food choices, and total heroin choices. These dependent measures were also analyzed using a repeated-measures two-way ANOVA (naltrexone) or mixed-effects analysis (lorcaserin) with treatment drug dose and dependent measure as the fixed main effects. Following a significant interaction, a Dunnett’s test was performed to compare treatment effects to baseline within a unit heroin dose or session dependent measure. Following a significant effect of treatment in the absence of an interaction, a Dunnett’s test was performed to compare collapsed treatment effects to baseline, irrespective of unit heroin dose or session dependent measure. The criterion for significance was set *a priori* at the 95% confidence level (*p*<0.05).

### 2.4 Drugs

Heroin HCl and (-)-naltrexone HCl were provided by the National Institute on Drug Abuse Drug Supply Program (Bethesda, MD) and dissolved in sterile saline. Lorcaserin HCl was purchased from a commercial supplier (MedChem Express, Monmouth Junction, NJ) and dissolved in sterile water. Heroin, naltrexone, and lorcaserin solutions were passed through a 0.22-micron sterile filter (Millex GV, Millipore Sigma, Burlington, MA) before administration. All drug doses were expressed as the salt forms listed above and delivered mg/kg based on weights collected bi-weekly to monthly.

## 3.0 Results

### 3.1 Baseline heroin vs. food choice and effect of saline substitution

Under baseline conditions, monkeys primarily responded on the food-associated key when heroin was not available (0 mg/kg/injection) or the unit heroin dose was small (0.001 mg/kg/injection) and almost exclusively reallocated their behavior to heroin choice during availability of larger unit heroin doses (0.01-0.032 mg/kg/injection) (**Figures 1A, 2A, 3A**; dashed lines). In addition, monkeys generally completed each of the ten possible ratio requirements of each component, although rate-decreasing effects of 0.032 mg/kg/injection heroin were observed (**Figures 1B, 2C, 3C**; dashed lines). **Figure 1** shows the average of the last three days of a 7-day saline substitution for heroin on percent choice (A) and the number of choices completed per component (B). Saline substitution resulted in a significant behavioral reallocation away from previously reinforced heroin-associated key and associated discriminative stimuli towards the food-associated key during components associated with previous 0.0032-0.032 mg/kg/injection heroin availability (**Figure 1A**: choice component: F_1.1,5.5_ = 37.9, *p* = 0.001; experimental manipulation: F_1,5_ = 48.3, *p* = 0.001; interaction: F_1.3,6.3_ = 31.2, *p* = 0.001). Furthermore, monkeys increased the number of completed choices per component under saline substitution conditions (**Figure 1B**: choice component: F_1.1,5.2_ = 154.4, *p* < 0.0001; experimental manipulation: F_1,5_ = 93.1, *p* = 0.0002; interaction: F_1.1,5.2_ = 154.4, *p* < 0.0001).

**Figure 1.**
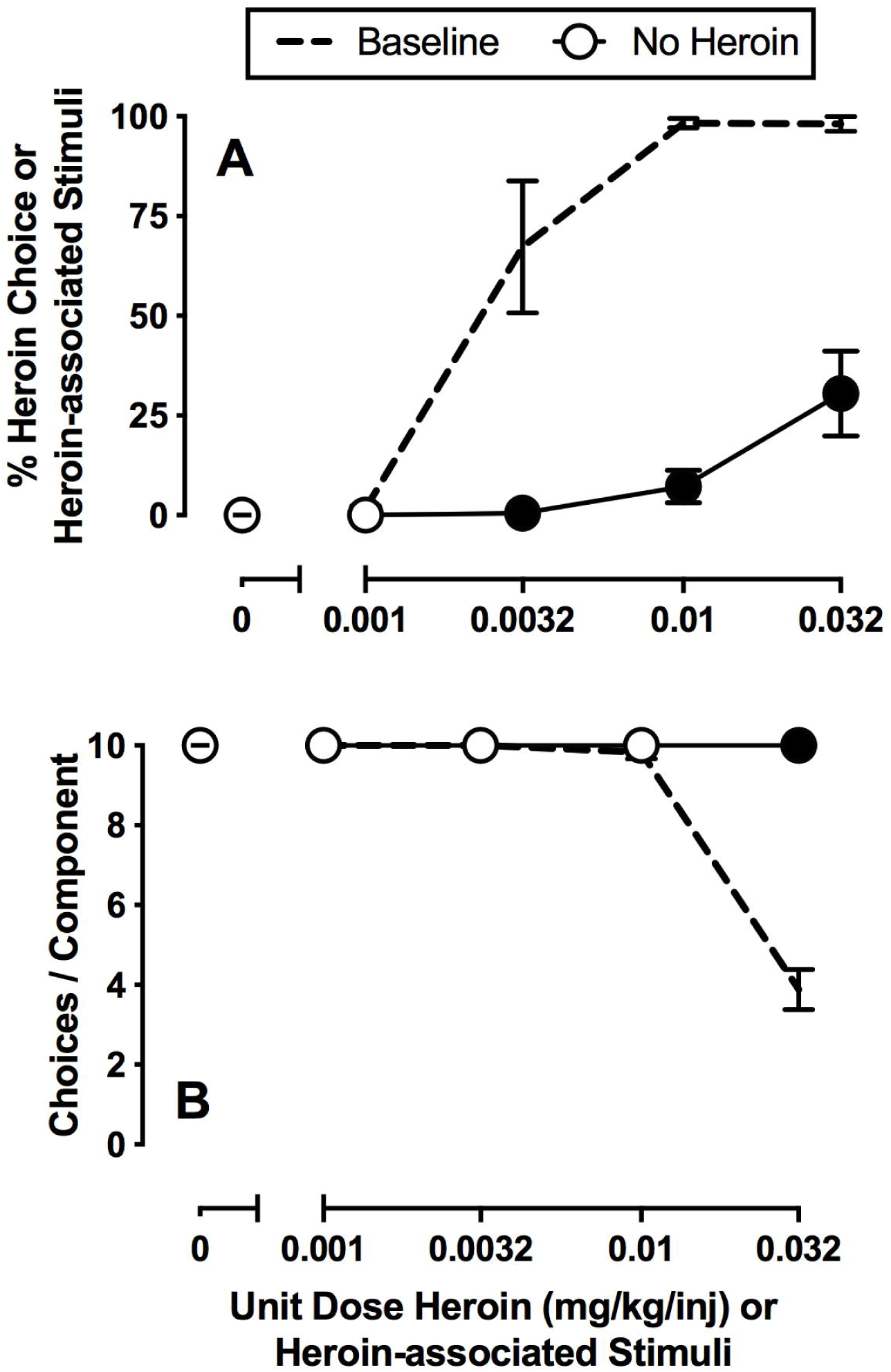
Effects of 7-day saline substitution on choice between heroin and food in female rhesus monkeys (*n*=6). Top and bottom abscissae: unit dose of heroin in milligrams per kilogram per injection or heroin-associated stimuli. Top ordinate (A): percent choice of heroin or heroin-associated stimuli. Bottom ordinate (B): number of choices completed per component. Points represent mean ±SEM obtained during days 5-7 of each 7-day treatment period. Filled symbols denote significantly different from baseline (*p*<0.05).

**Figure 2.**
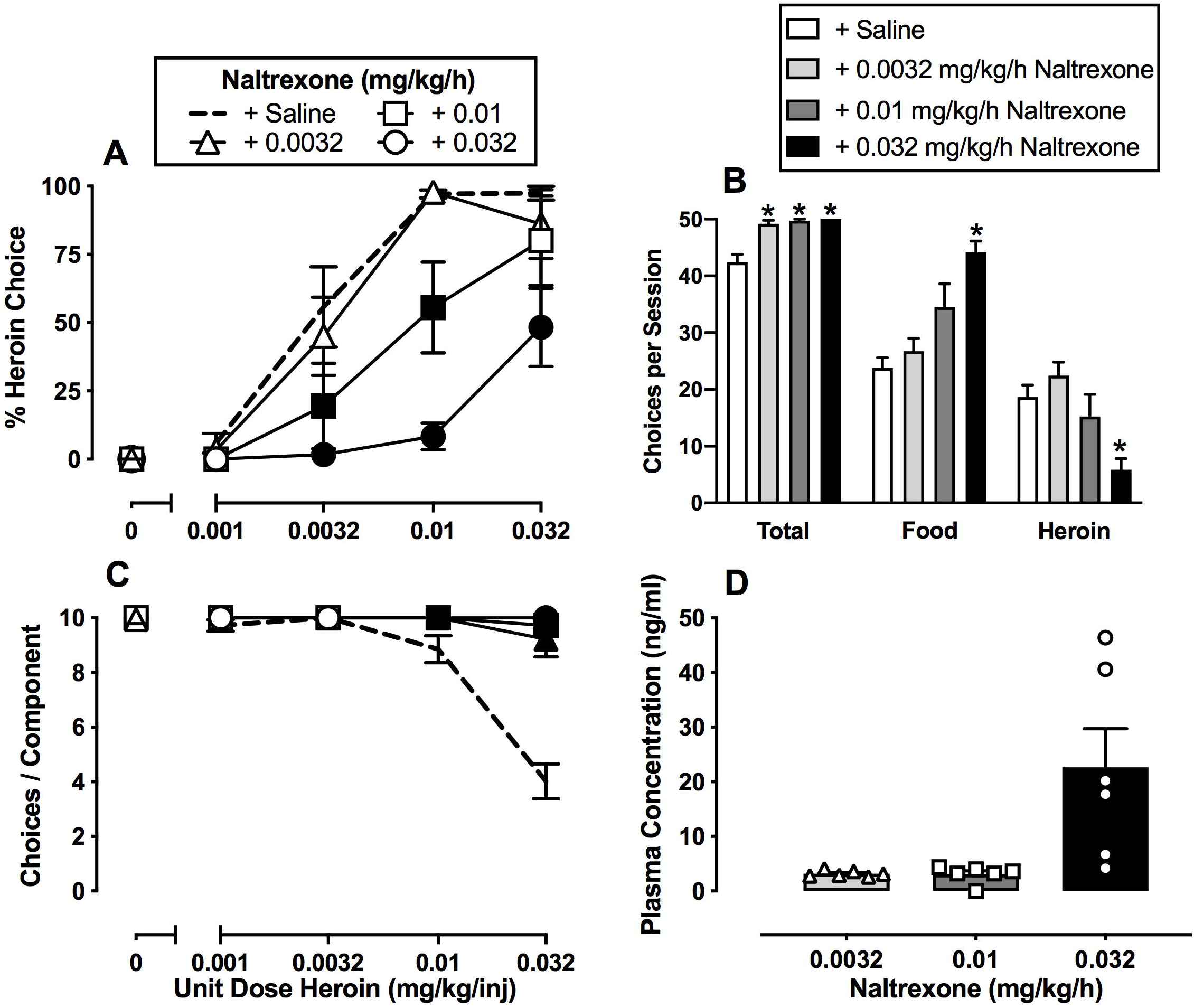
Effects of 7-day intravenous naltrexone (0.0032-0.032 mg/kg/h) treatment on choice between heroin and food in female rhesus monkeys (*n*=6). Left abscissae: unit dose of heroin in milligrams per kilogram per injection. Top left ordinate (A): percent heroin choice. Bottom left ordinate (C): number of choices completed per component. Top right abscissae: dependent measure. Top right ordinate (B): number of choices made per session. Bottom right abscissae: dose of naltrexone (mg/kg/h). Bottom right ordinate (D): plasma naltrexone concentration (ng/ml). Points and bars of A, B, and C represent mean ±SEM obtained during days 5-7 of each 7-day treatment period. Bars of D represent mean ±SEM. Filled symbols in the left panels or * in the top right panel denote significantly different from saline treatment (*p*<0.05).

**Figure 3.**
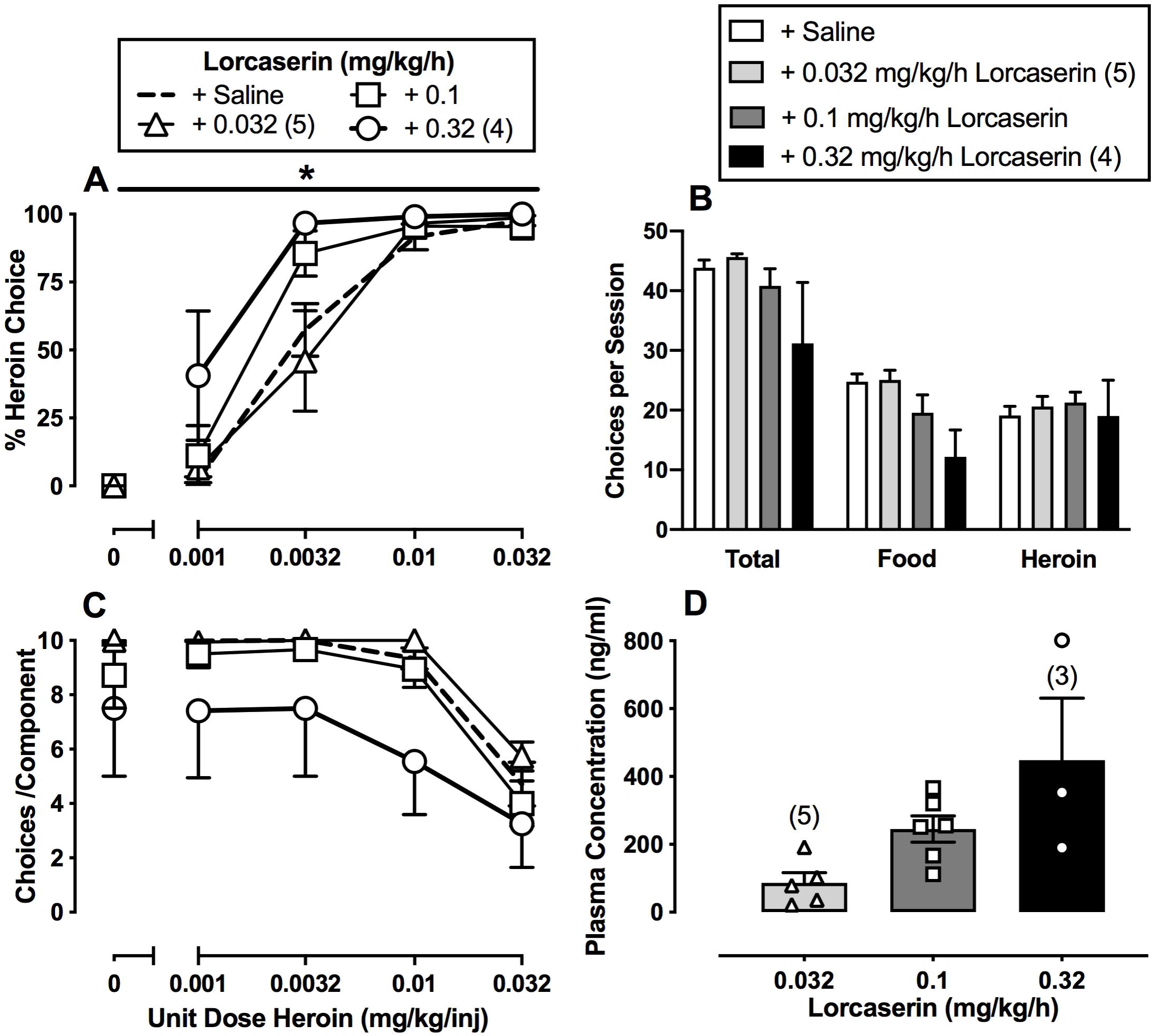
Effects of 7-day intravenous lorcaserin (0.032-0.32 mg/kg/h) treatment on choice between heroin and food in male (*n*=1) and female (*n*=5) rhesus monkeys. Left abscissae: unit dose of heroin in milligrams per kilogram per injection. Top left ordinate (A): percent heroin choice. Bottom left ordinate (C): number of choices completed per component. Top right abscissae: dependent measure. Top right ordinate (B): number of choices made per session. Bottom right abscissae: dose of lorcaserin (mg/kg/h). Bottom right ordinate (D): plasma concentration (ng/ml) of lorcaserin. Points and bars of A, B, and C represent mean ±SEM obtained during days 5-7 of each 7-day treatment period. Bars of D represent mean ±SEM. Numbers in parentheses indicate the number of monkeys contributing to that data point if fewer than the total sample size (*n*=6). * denotes collapsed 0.32 mg/kg/h lorcaserin-treatment data are significantly (*p*<0.05) different from saline.

### 3.2 Effect of naltrexone on heroin choice

**Figure 2** shows the average of the last three days of 7-day continuous naltrexone treatment (0.0032-0.032 mg/kg/h) on heroin-vs.-food choice. Naltrexone dose-dependently decreased heroin choices and reciprocally increased food choices (**Figure 2A**: heroin dose: F_2.4,12_ = 79.6, *p* < 0.0001; naltrexone dose: F_1.5,7.6_ = 10.8, *p* = 0.008; interaction: F_2.6,13.2_ = 4.8, *p* = 0.02). In addition, naltrexone treatment increased the number of choices completed per component (**Figure 2C**: heroin dose: F_1.1,5.2_ = 154.4, *p* < 0.0001; naltrexone dose: F_1,5_ = 93.1, *p* = 0.0002; interaction: F_1.1,5.2_ = 154.4, *p* < 0.0001). All three naltrexone doses significantly increased total session choices and 0.032 mg/kg/h naltrexone significantly increased total food choices and decreased total heroin choices (**Figure 2B**: dependent measure: F_1,5.1_ = 144.5, *p* < 0.0001; naltrexone dose: F_1.3,6.7_ = 18.8, *p* = 0.0028; interaction: F_1.6,7.8_ = 8.8, *p* = 0.0125). As shown in **Figure 2D**, seven days of 0.032 mg/kg/h naltrexone treatment resulted in plasma naltrexone concentrations of approximately 20 ng/ml. Notably, the initiation of naltrexone treatment did not precipitate overt somatic withdrawal signs (i.e., diarrhea, wet dog shakes, or lying on the bottom of the cage) in any monkey during informal evaluation throughout the day, suggesting an absence of opioid dependence.

### 3.4 Effect of lorcaserin on heroin choice

**Figure 3** shows the average of the last three days of seven days of continuous lorcaserin treatment (0.032-0.32 mg/kg/h) on heroin-vs.-food choice. Lorcaserin (0.32 mg/kg/h) treatment significantly increased heroin-vs.-food choice compared to baseline (**Figure 3A**: heroin dose: F_1.6,7.8_ = 130.2, *p* < 0.0001; lorcaserin dose: F_2.5,12.6_ = 4.7, *p* = 0.02). Lorcaserin treatment did not significantly alter the number of choices completed per component (**Figure 3C**: heroin dose: F_1.1,5.7_ = 23.69, *p* < 0.01) or total session choices, food choices, or heroin choices (**Figure 3B**). However, the highest lorcaserin dose (0.32 mg/kg/h) produced overt signs of sedation (data not shown), and higher lorcaserin doses were not tested due to safety concerns. Across the dose range tested, lorcaserin produced dose-dependent increases in plasma lorcaserin levels, and 7-day 0.32 mg/kg/h lorcaserin treatment resulted in plasma concentrations of approximately 450 ng/ml (**Figure 3D**).

## 4.0 Discussion

The present study determined the effectiveness of repeated 7-day treatment with the clinically available 5-HT_2C_ agonist lorcaserin to attenuate heroin-vs.-food choice in rhesus monkeys. For comparison, 7-day naltrexone treatment and saline substitution for heroin were determined as positive controls. There were three main findings. First, saline substitution for heroin resulted in an attenuation, but not elimination, of responding on the previously heroin-associated key and a reciprocal increase in responding on the food-associated key. Second, 7-day naltrexone treatment significantly decreased heroin choice and produced a reciprocal increase in food choice similar to saline substitution and consistent with the clinical effectiveness of depot naltrexone for OUD. Lastly, 7-day lorcaserin treatment significantly increased heroin-vs.-food choice. Overall, the present results do not support the continued evaluation and clinical utility of lorcaserin as a candidate OUD medication.

### 4.1 Effects of saline substitution and naltrexone treatment

The present study compared the effects of three different treatments designed to reduce or eliminate the centrally mediated abuse-related effects of heroin. Specifically, saline substitution removed the possibility of heroin delivery and heroin or metabolites (i.e. 6-acetylmorphine or morphine) binding to MORs, whereas naltrexone allowed heroin delivery and distribution to MORs in brain but decreased the probability of MOR binding by heroin and its metabolites through competitive interactions at MORs. In contrast, lorcaserin allowed heroin delivery, distribution, and heroin or its metabolites binding to MORs but is intended to oppose heroin-induced activation of brain reward systems. The first of these approaches, saline substitution, eliminated heroin from the environment and shows the maximum effects that can be achieved with antagonist approaches intended to block heroin effects. The elimination of heroin from the environment is the principal goal of “supply side” anti-drug interventions ((Greenfield and Paoli, 2017), e.g. policies to prevent the manufacturing and distribution of heroin)), and it is the persistence of heroin availability despite supply-side interventions that necessitates development of effective OUD treatments for vulnerable individuals. Nonetheless, the present results with saline substitution provide evidence to suggest that that blocking heroin effects has the potential to reduce heroin-maintained behavior and promote more adaptive behaviors maintained by non-drug reinforcers. However, approximately 30% responding on the heroin-associated key was observed at the end of 7-day saline substitution experiment and mostly during the last component when the largest heroin dose was previously available. This result suggests that some procedural aspect(s) (e.g., visual discriminative stimuli, temporal characteristics) might function as conditioned stimuli that maintain behavior in the absence of heroin delivery that has not extinguished within 7 days. This finding should be considered when interpreting the effects of pharmacological treatments described below.

This desirable outcome of behavioral allocation away from heroin and towards non-drug reinforcers (i.e. food) could also be achieved with appropriate doses of the FDA-approved maintenance medication naltrexone. A naltrexone dose of 0.032 mg/kg/h was necessary to produce a minimal 8-fold potency shift in the heroin-vs.-food choice dose effect function and resulted in plasma naltrexone levels of approximately 20 ng/mL. Plasma naltrexone levels of ≥2 ng/mL in humans following depot naltrexone administration are sufficient to antagonize effects of 25mg IV heroin (Comer et al., 2002; Sullivan et al., 2006). For comparison, oral naltrexone 50 mg results in peak naltrexone levels of approximately 10 ng/mL in humans (Dunbar et al., 2006). The present naltrexone results are consistent with previous acute and continuous naltrexone treatments on opioid-vs.-food choice in rats (Townsend et al., 2019) and monkeys (Maguire et al., 2019), and continuous naloxone treatment effects on heroin-vs.-food choice in monkeys (Negus, 2006). Furthermore, the present results are also consistent with depot naltrexone treatment effects on heroin-vs.-money choice in humans (Comer et al., 2006) and clinical trials examining naltrexone effectiveness (Comer et al., 2006; Krupitsky et al., 2011; Lee et al., 2018). Overall, these naltrexone results provide an empirical framework for interpreting the effectiveness of emerging candidate OUD medications such as lorcaserin.

### 4.2 Lorcaserin effects on heroin choice

Repeated lorcaserin treatment failed to attenuate heroin-vs.-food choice in monkeys and, in fact, significantly increased heroin-vs.-food choice. Peak lorcaserin plasma levels during the present study were approximately 5-fold larger than peak lorcaserin levels reported in humans after treatment with clinically available formulations used for obesity treatment (Christopher et al., 2016), suggesting that a sufficient dose range was examined. Moreover, the highest dose tested in the present study produced overt signs of sedation in some monkeys, and higher doses were not tested due to safety concerns. The present lorcaserin results on heroin-vs.-food choice in monkeys are consistent with a previous remifentanil-vs.-food choice study in rats wherein acute lorcaserin pretreatment was found to produce non-selective decreases in rates of self-administration without significantly affecting behavioral allocation between the two reinforcers (Panlilio et al., 2017). Previous studies in rats (Neelakantan et al., 2017) and nonhuman primates (Gerak et al., 2019; Kohut and Bergman, 2018) found that lorcaserin dose-dependently decreased opioid self-administration and opioid-induced reinstatement of extinguished opioid self-administration after acute treatment. Furthermore, decreases in opioid self-administration were sustained during repeated lorcaserin treatment, and both acute and repeated lorcaserin-treatment effects displayed some degree of behavioral selectivity insofar as opioid self-administration was decreased without a significant decrease in food-maintained responding (Kohut and Bergman, 2018). Given that a range of doses spanning ineffective lorcaserin doses (i.e., 0.032 mg/kg/h) to a dose (i.e., 0.32 mg/kg/h) that significantly increased choice of heroin over food, we hypothesize differences in lorcaserin effectiveness to selectively attenuate opioid-vs. food-maintained behavior were not due to differences in route of lorcaserin administration (e.g., repeated bolus dosing versus continuous infusion), but rather behavioral procedure differences in how opioid reinforcement was measured (i.e., rates of self-administration in the previous studies compared to behavioral allocation between drug and food in the present study). Although behavioral selectivity to decrease opioid self-administration is a desirable attribute of candidate medications (Banks et al., 2019; Mello and Negus, 1996), the expression of behavioral selectivity can be influenced by experimental variables independent of the candidate medication. The present results suggest the degree of behavioral selectivity shown with lorcaserin is not sufficient to effectively decreases opioid reinforcement.

One of these experimental variables is the unit heroin dose available during the self-administration session. For example, repeated 1 mg/kg/day lorcaserin decreased rates of 3.2 and 10 μg/kg/injection heroin self-administration, but not food-maintained responding, in rhesus monkeys when assessed under a multi-day progressive-ratio procedure (Kohut and Bergman, 2018). Consistent with these previous results, there was a non-significant trend in the present study for 0.32 mg/kg/h lorcaserin to decrease the number of choices completed per component during 10 μg/kg/injection unit heroin dose availability and during a choice component where heroin-vs.-food choice was 100% under baseline conditions. However, unlike naltrexone treatment effects in Figure 2, lorcaserin failed to promote behavioral reallocation away from heroin and towards the alternative non-drug food reinforcer at any unit heroin dose. OUD medications that decrease both large- and small-dose heroin self-administration should be prioritized because OUD typically involves consumption of large drug doses and because behavior maintained by small drug doses are often sensitive to nonpharmacological strategies such as contingency management or the therapeutic workplace (Jarvis et al., 2017).

Lorcaserin and other 5-HT_2c_ agonists are under consideration as candidate OUD medications based on evidence that they might be effective to oppose and limit abuse-related MOR agonist activation of the mesolimbic DA system. However, the present results suggest that this strategy for functional antagonism of MOR agonist abuse-related effects is not as effective as direct MOR antagonism with naltrexone. Rather, these results suggest that sustained 5-HT_2c_ receptor activation may produce both direct effects that pose safety concerns and a basal tone of neural activity that actually increases the relative reinforcing efficacy of MOR agonists in comparison to food as a non-drug alternative reinforcer.

### 4.3 Conclusions

In conclusion, the present lorcaserin treatment effects on heroin-vs.-food choice parallel differences in lorcaserin treatment effects on cocaine self-administration based on the schedule of reinforcement. For example, repeated lorcaserin treatment decreased rates of cocaine self-administration under fixed-ratio and progressive-ratio schedules of reinforcement (Collins et al., 2016; Gerak et al., 2016), but failed to attenuate cocaine-vs.-food choice under a concurrent schedule of reinforcement (Banks and Negus, 2017a) in rhesus monkeys (for review, see (Collins et al., 2017)). Recently, lorcaserin failed to attenuate cocaine-vs.-money choice and enhanced subjective effects of cocaine in humans (Pirtle et al., 2019). Although there are currently no published human laboratory studies or clinical trials examining lorcaserin treatment effects on opioid-taking behavior, results from an unpublished human laboratory study found lorcaserin maintenance also failed to attenuate oxycodone-vs.-money choice (Comer, personal communication). Consistent with the present results, these human laboratory results provide further evidence against the clinical utility of lorcaserin as a candidate OUD medication. Moreover, the present results further highlight the utility of preclinical drug-vs.-food choice procedures to efficiently evaluate candidate OUD medications to prioritize novel therapeutics for promotion to human laboratory drug self-administration studies and clinical trials (Banks et al., 2019).

## Contributors

EAT, SSN, and MLB were responsible for study concept, design, and execution. EAT and MLB drafted the initial manuscript. JLP performed the sample preparation and analysis of the plasma naltrexone and lorcaserin levels. All authors critically reviewed the manuscript for content and approved the final manuscript version submitted for publication. We appreciate the technical assistance of Jennifer Gough, Kaycee Faunce, and Rebekah Tenney. We also acknowledge Kevin Costa for writing the original version of the behavioral program that was modified for the choice studies conducted in this manuscript.

